# Two sides of the same coin: monetary incentives concurrently improve and bias confidence judgments

**DOI:** 10.1101/099382

**Authors:** Maël Lebreton, Shari Langdon, Matthijs J. Slieker, Jip S. Nooitgedacht, Anna E. Goudriaan, Damiaan Denys, Ruth J. van Holst, Judy Luigjes

**Affiliations:** Amsterdam Brain and Cognition (ABC); CREED, Amsterdam School of Economics (ASE), Universiteit van Amsterdam, 1018 WB Amsterdam, the Netherlands.; Amsterdam Institute for Addiction Research; Department of Psychiatry, Academic Medical Centre, 1100 DD Amsterdam, the Netherlands; Netherlands Institute for Neuroscience, Institute of the Royal Netherlands Academy of Arts and Sciences, 1105 BA Amsterdam, the Netherlands

## Abstract

Decisions are accompanied by a feeling of confidence, i.e., a belief about the decision being correct. Confidence accuracy is critical, notably in high-stakes situations such as medical or financial decisionmaking. Here, we investigated how incentive motivation influences confidence accuracy by combining a perceptual task with a confidence incentivization mechanism. Importantly, by varying the magnitude and valence (gains or losses) of monetary incentives, we orthogonalized their motivational and affective components. Corroborating theories of rational decision-making and motivation, our results first reveal that the motivational value of incentives improves aspects of confidence accuracy. However, in line with a value-confidence interaction hypothesis we further show that the affective value of incentives concurrently biases confidence reports, thus degrading confidence accuracy. Finally, we demonstrate that the motivational and affective effects of incentives differentially impact how confidence builds on perceptual evidence. Altogether, these findings may provide new hints about confidence miscalibration in healthy or pathological contexts.

## Introduction

Imagine you have to cross a road. It's dark and raining, the visibility is low. At some point, you estimate that there does not seem to be any danger, and decide to cross. However, because you feel quite unsure about crossing with such low visibility, you decide to check one last time, and spot a car coming at high speed to your direction. Luckily, you have enough time to withdraw from the street and avoid being hit by the car.

Just like in this example, most decisions in everyday life are accompanied by a subjective feeling of confidence emerging from the constant monitoring of our own thoughts and actions by metacognitive processes (Fleming and Dolan, 2012; Yeung and Summerfield, 2012). Formally, confidence is a decision maker's estimate of the probability – or belief - that her action, answer or statement is correct, based on the available evidence (Adams, 1957; Pouget et al., 2016). Measuring confidence accuracy – i.e. the quality of metacognitive judgments - is challenging (Brier, 1950; Fleming and Lau, 2014; Nelson, 1984), but confidence accuracy can consensually be split in a *bias* (or calibration) component measuring how confidence judgments differ from the overall probability of being correct, and a *sensitivity* (or discrimination) component measuring how reliably confidence judgments can dissociate correct from incorrect answers (Brier, 1950; Fleming and Lau, 2014).

Although high confidence accuracy seems critical to monitor and re-evaluate previous decisions (Folke et al., 2016), to track changes in the environment (Meyniel et al., 2015a), or to arbitrate between different strategies (Donoso et al., 2014; Vinckier et al., 2016), converging evidence suggests that confidence judgments are significantly *biased*. Notably, we often overestimate the probability of being correct, a phenomenon called overconfidence (Lichtenstein et al., 1982). This bias, potentially detrimental for the decision-maker or society, has been consistently reported in numerous domains and situations, from simple sensory psychophysics (Baranski and Petrusic, 1994) or knowledge (West and Stanovich, 1997) tasks in the laboratory, to medical (Berner and Graber, 2008), financial, and managerial (Camerer and Lovallo, 1999; Malmendier and Tate, 2005) decision-making.

Altogether, the importance of confidence as a cognitive variable mitigating decision-making and the societal relevance of its miscalibration have considerably stimulated the search for factors which modulate or bias confidence accuracy. An influential line of research, encapsulated under the term “*motivated cognition*” or “*motivated reasoning*”, has suggested that beliefs are influenced by individuals' desires (Bénabou and Tirole, 2016; Epley and Gilovich, 2016; Kunda, 1990). In other terms, individuals tend to estimate desirable events to be more likely than undesirable ones, potentially leading to overconfidence (Giardini et al., 2008). In a different line of research, studies have established links between incidental psychological states such as elevated mood (Koellinger and Treffers, 2015), absence of worry (Massoni, 2014) or emotional arousal (Allen et al., 2016; Jönsson et al., 2005) and (over-)confidence. Recently, functional neuroimaging studies reported neural correlates of confidence in the ventromedial prefrontal cortex (De Martino et al., 2013; Lebreton et al., 2015), as well as in mesolimbic and striatal regions (Guggenmos et al., 2016; Hebart et al., 2014), a brain network associated with the encoding of economic, motivational and affective values (Levy and Glimcher, 2012). Such an overlap in the neural correlates of confidence and values suggests that these variables also interact at the behavioral level. In practice, this hypothesis entails that a decision-maker reports higher confidence not only because she believes she is correct, but also because she is in a high expected - or experienced - value context. This value-confidence interaction could both parsimoniously explain associations between positive affective-states and overconfidence (Allen et al., 2016; Giardini et al., 2008; Jönsson et al., 2005; Koellinger and Treffers, 2015; Massoni, 2014), and underpin biases in confidence judgments.

In this report, we methodically investigated the interactions between incentive motivation and confidence, in an attempt to explain features of human confidence accuracy. To do so, we designed a task where participants had to first make a difficult perceptual decision, and then judge the probability of their answer being correct, i.e., their confidence in their decision (**Fig. 1.A**). In order to identify the critical features of the interactions between incentive motivation and confidence accuracy, the accuracy of confidence was incentivized with monetary prospects whose magnitude and valence were systematically varied (see **Fig. 1.** and **Methods**). This experimental manipulation elegantly orthogonalized the net incentive value (i.e. the affective component of incentives, which can take both positive and negative values, and thereafter indexed as V) and the absolute incentive value (i.e. the motivational value of incentives, regardless of their valence, thereafter indexed as |V|). We used this experimental setup to investigate the effects those two aspects of incentive on the two core components of confidence accuracy: bias and sensitivity.

**Fig. 1.**
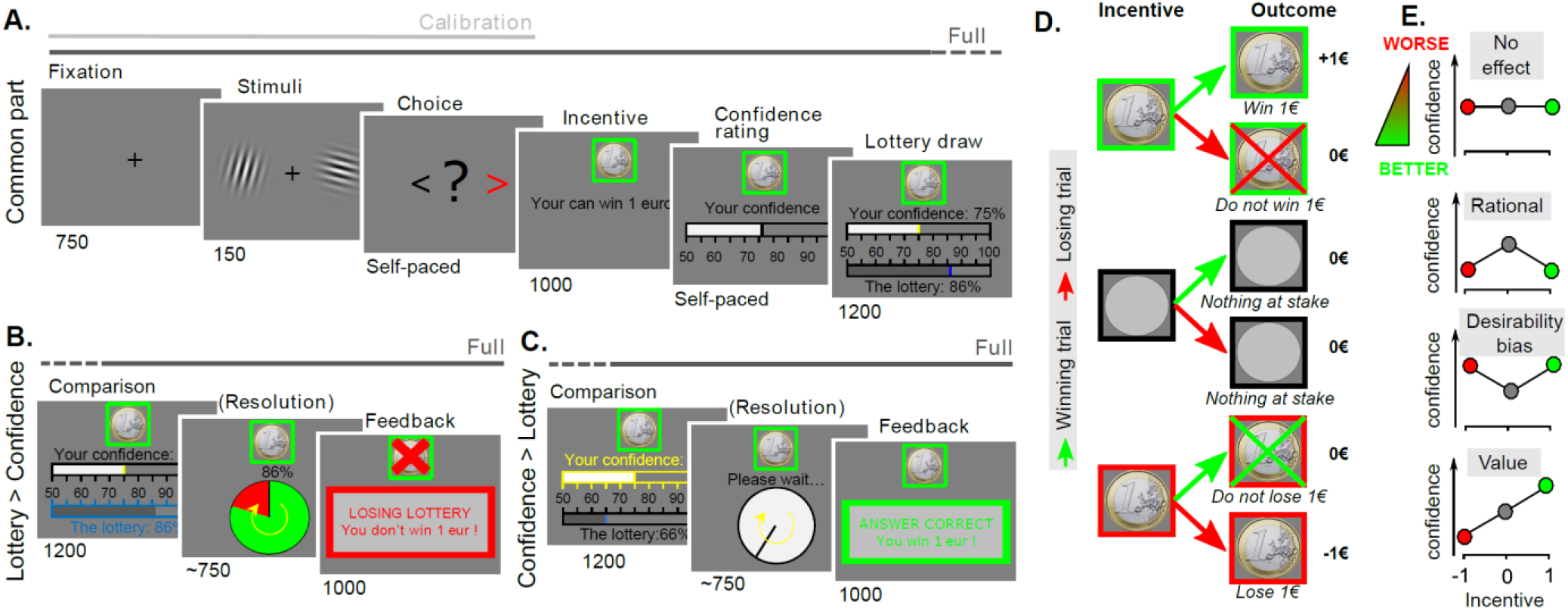
Behavioral task and hypotheses. Successive screens displayed in one trial are shown from left to right with durations in ms. **A. Behavioral task - Common part**. Participants viewed a couple of Gabor patches displayed on both sides of a computer screen, and judged which had the highest contrast. They were thereafter presented with a monetary stake (in a green frame for gain, grey for neutral and red for losses), and asked to report their confidence C in their answer on a scale from 50 to 100%. Then, a lottery number L was drawn in a uniform distribution between 50 and 100%, displayed as a scale under the confidence scale and the scale with the highest number was highlighted. **B. Behavioral task - Lottery > Confidence.** If the L > C, the lottery was implemented. A wheel of fortune, with a L% chance of losing was displayed, and played. Then, *feedback* informed whether the lottery resulted in a win or a loss. **C. Behavioral task - Confidence > Lottery.** If C > L, a clock was displayed together with the message “*Please wait*”, followed by *feedback* which depended on the correctness of the initial choice. Subject would win (gain frame) or not lose (loss frame) the incentive in case of a “winning” trial, and they would not win (gain frame) or lose (loss frame) the incentive in case of a “losing” trial. **D. Behavioral task – Payoff matrix.** Depending on the combination of a trial's offered incentive and the trial's final win or loss (regardless of whether the lottery or the correctness of the answer determined it), participants could receive various outcomes, from winning the proposed incentive to losing the proposed incentive. **E. Hypotheses.** The expected biasing effects of incentives (-1€; 0€ or +1€) on confidence, under different theoretical hypotheses. (1^st^ line) H0: No biasing effects of incentives. Participants are similarly overconfident in the 3 incentive conditions. (2^nd^ line) H1: rational decision making. Under higher incentives, participants are more rational, i.e. less biased. The absolute value of incentives therefore decreases confidence, if participants are generally overconfident. (3^rd^ line) H2: desirability bias. Participants are more inclined to believe that they are correct when higher incentives are at stake. The absolute value of incentives increases confidence. (4^th^ line) H3: value-confidence interaction. The confidence judgment of participants is affected by the affective component of incentives. The incentive net value impacts confidence.

Standard theories of rational decision-making and motivation from behavioral economics (Bonner and Sprinkle, 2002; Smith and Walker, 1993; Wilcox, 1993) and cognitive psychology (Botvinick and Braver, 2015) predict that higher stakes increase participants tendency to conform to rational model predictions, which should improve confidence accuracy to maximize the payoff (as incentivized by the MP mechanism, see **Methods** and **SI results**). Higher absolute incentive value should therefore increase confidence *sensitivity,* irrespective of the incentive valence (gain or loss).

Besides, orthogonalizing the affective and motivational components of incentives enabled us to test three opposing predictions from three different theories anticipating effects of incentives on confidence *bias* (**Fig. 1.E**). First, as outlined in the previous paragraph, standard theories of rational decision-making predict that higher stakes improve confidence accuracy and therefore also a reduced metacognitive bias is expected. In this case, we expect that if participants are generally biased toward overconfidence, an increase in absolute incentive value should reduce this bias, and therefore decrease confidence judgments. Second, motivated cognition theories (Kunda, 1990) – i.e. in the form of the desirability bias (Giardini et al., 2008)- predict that participants should be more motivated to believe that they are correct when more money is at stake, irrespective of the valence (gain or loss). In this case, an increase in absolute incentive value should increase confidence judgments (and exaggerate the overconfidence bias). Finally, our value - confidence interaction hypothesis predicts higher monetary incentives should bias confidence judgments upwards in a gain frame, and downward in a loss frame, despite the potentially detrimental consequences on the final payoff. In this case, the net incentive value should bias confidence judgments.

In four experiments, we repeatedly found behavioral patterns which confirm the motivational effect of incentives on confidence sensitivity, but pinpoint biasing effect of incentives in line with value-confidence interaction hypothesis.

## Results

We collected data in four experiments in which participants performed different versions of a confidence elicitation task (**Fig. 1**, **Table 1** and **Methods**); in each trial, participants briefly saw a pair of Gabor patches first, then had to indicate which one had the highest contrast, and finally had to indicate how confident they were in their answer (from 50 to 100%). Critically, the confidence judgment was incentivized: after the binary choice and before the confidence judgment, a monetary stake was displayed, which could be neutral (no incentive) or indicate the possibility to gain or lose a certain payoff (e.g. 10 cents, 1 euro, 2 euros), which differed between the experiments. Participants could maximize their chance to gain (respectively not lose) the stake by reporting their confidence as accurately and truthfully as possible, because the outcome of trial was determined by a Matching Probability (MP) mechanism, a well-validated method from behavioral economics adapted from the Becker-DeGroot-Marschak auction (Becker et al., 1964; Ducharme and Donnell, 1973). Briefly, the MP mechanism considers participants' confidence reports as bets on the correctness of their answers, and implements trial-by-trial comparisons between these bets and random lotteries. Under utility maximization assumptions, this guarantees that participants maximize their earnings by reporting their most precise and truthful confidence estimation (Schlag et al., 2015; Schotter and Trevino, 2014). Because this incentivization was implemented after the perceptual choice, it is possible to separately motivate the accuracy of confidence judgments without directly influencing the performance on the perceptual decision. Prior to the task, participants performed a calibration session, which was used to generate the main task stimuli, such that the decision situations spanned a pre-defined range of individual subjective difficulties (see **Methods**).

**Table.1:** Demographics and experimental design. P(c=L) indicates the level of difficulty (i.e probability of choosing the left Gabor) used to generate the stimuli. The number of times all levels of difficulty were repeated per incentive and per session is indicated between brackets. Tasks indicate which task version (short: confidence incentivization Short Version, extended: confidence incentivization Extended Version, perf: Performance incentivization Version) was offered to participants, and the number of session per task is indicated between brackets.

|  | Exp. 1 | Exp. 2 | Exp. 3 | Exp. 4 |
| --- | --- | --- | --- | --- |
| M/F (total) | 6/18 (24) | 7/17 (24) | 9/26 (35) | 7/17 (24) |
| Age | 23.8±6.00 | 23.0±5.67 | 24.8±5.43 | 24.3±7.79 |
| Stakes (€) | 0; ±1; | 0; ±1; | ±0.1; ±1; | ±0.1; ±1;±2 |
| P(c=L) | 50 ± [10:10:40] (×4) | 50±[15:10:35] (×2) | 50±[0:05:15] (×2) | 50±[0:10:40] (×2) |
| Tasks | Short (×4) | Short (×2) + Perf (×2) | Ext. (×1) + Short(×1) | Short (×3) |

### Basic features of confidence judgments

As a pre-requisite, we assessed the quality of our experimental design and the validity of our experimental variables, irrespective of the effects of monetary incentives. We notably show that, in all four experiments, *ex ante* choice predictions from our psychophysical model closely match participants' actual choice behavior (**SI results**). Additionally, we show that in all four experiments, participants' confidence judgments exhibit three fundamental properties (Sanders et al., 2016): (1) confidence ratings correlate with the probability of being correct; (2) the link between confidence ratings and evidence is positive for correct and negative for incorrect responses; (3) the link between evidence and performance differs between high and low confidence trials (**SI results**). Overall, these preliminary results suggest that the confidence measure elicited in our task actually corresponds to subjects' estimated posterior probability of being correct (Fleming and Daw, 2017; Sanders et al., 2016). They also address potential concerns about the validity of confidence elicitation in general (Drugowitsch, 2016; Drugowitsch et al., 2014), and additionally demonstrate that our MP incentivization mechanism did not bias or distort confidence.

### Effects of incentives on confidence judgments

Twenty-four subjects participated in our first experiment, where the combination of their choice and confidence ratings could lead, depending on the trial, to a gain or no-gain of 1 euro, a loss or no-loss of 1 euro, or to a neutral outcome (**Fig. 1.C**). In order to investigate the interaction between incentive motivation and confidence, and compare the predictions of the different theories (**Fig 2.A**), we implemented linear mixed-effect models, with the net and absolute incentive values as independent variables (see **Methods**). In line with our value-confidence interaction hypothesis, our results first show that participants' confidence judgments are specifically modulated by the incentive net value (β_V_ = 2.06±0.42, p<0.001; β_|V|_ = -0.97±1.03, p=0.38). Critically, and as expected from our task design, this effect of incentives on confidence is not driven by an effect on performance, given that neither the net nor absolute incentive value have any effect on performance (both p>0.20).

### Effects of incentives on confidence (metacognitive) accuracy

In order to explore how incentives impact confidence *accuracy*, we adopted the signal-detection (SDT) approach developed for metacognition (Fleming and Lau, 2014; Maniscalco and Lau, 2012; Massoni et al., 2014). STD postulates that both the binary choice and the metacognitive (confidence) estimation are based on the same noisy source of perceptual evidence. The goal of SDT analysis is to estimate from the observed distributions of choices and confidence ratings how this internal signal is used by participants to derive their decisions. Under a few assumptions, the SDT framework can be used to dissociate and measure two components of metacognitive accuracy: the metacognitive bias, and the metacognitive sensitivity.

The metacognitive *bias* is the tendency to give high confidence ratings, all else being equal (Fleming and Lau, 2014). We used, as a measure of this bias, a classical measure of overconfidence (Hollard et al., 2015; Lichtenstein et al., 1982), computed as the difference between the averaged confidence and the averaged performance. Therefore, a metacognitive bias of zero signals high confidence accuracy, whereas a positive (respectively negative) calibration signals overconfidence (respectively underconfidence) and thus a lower confidence accuracy.

The metacognitive *sensitivity* measures the efficacy with which observers' confidence ratings discriminate between their own correct and incorrect answers. We used, as a metric for this sensitivity, the meta-d', which estimates how much information, in signal-to-noise (d) units, is available for the confidence judgment (Maniscalco and Lau, 2012). Therefore, the higher the meta-d’, the more sensitive confidence judgments are to the correctness of the choice. Critically, as opposed to most other metrics of confidence accuracy, the meta-d’ is not influenced by the response bias (such as average confidence level) or the ability to distinguish stimuli (i.e. to perform the binary choice task) (Fleming and Lau, 2014; Galvin et al., 2003; Maniscalco and Lau, 2012). Note however that, as expected from our task design (i.e. the incentives being uncovered *after* the binary choice), the SDT signal-to-noise metric quantifying this ability to distinguish stimuli (i.e. the d’) is not impacted by the incentive level in any of the tasks (see **SI results**).

Our results first show that metacognitive sensitivity (meta-d’) is specifically modulated by the absolute incentive value (β_V_ = β = 0.00 ± 0.05, p = 0.94; β_|V|_ = 0.34 ± 0.09, p< 0.001; **Fig. 2.B**); this means that positive and negative incentivization symmetrically improve participants' metacognitive sensitivity compared to the no-incentives condition. We refer to this first effect as the *motivational* effect of incentives on confidence accuracy. Second, mirroring the effects on confidence judgements, metacognitive bias monotonically increases with the net incentive value (β_V_ = 2.78±0.67, p < 0.001; β_|V|_ = -0.71±1.42, p = 0.62; **Fig 2.B**). Since participants are overconfident on average, metacognitive bias is thereby improved by loss prospects, but paradoxically deteriorated by gain prospects. We refer to this second effect as the *biasing* effect of incentives on confidence.

**Fig. 2.**
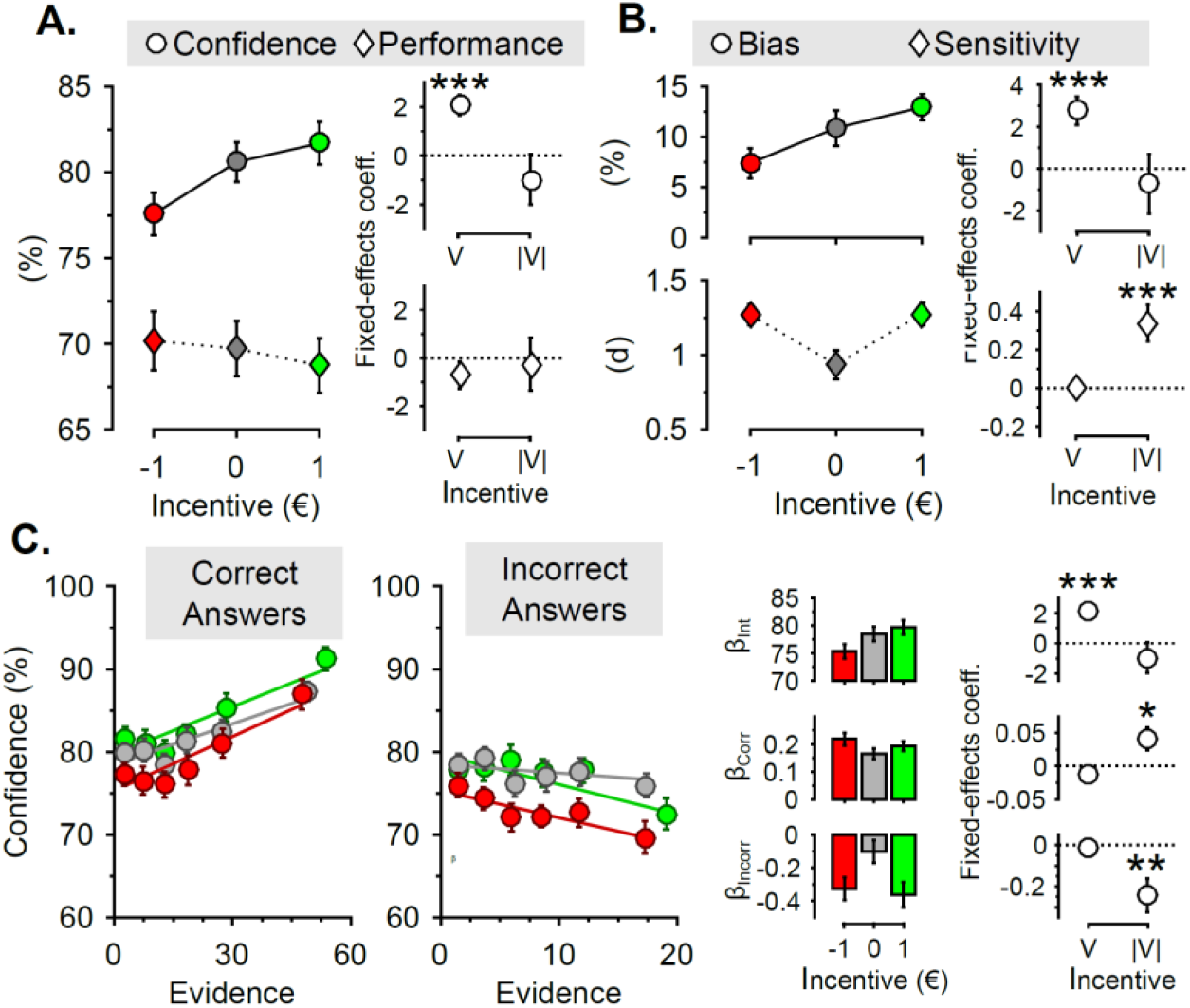
Experiment 1. **A. Incentive effects on behavior.** Reported confidence (dots) and performance (diamonds) – i.e. % correct-as a function of incentives. **B. Incentive effects on confidence (metacognitive) accuracy.** Computed bias (top, dots) and meta-d’ (diamonds) as a function of incentives. The insets presented on the right-hand side of the graphs (**A-B**) depict the results of the linear mixed-effect model, estimated for each behavioral (**A**. top: confidence; bottom: performance) and metacognitive (**B.** top: bias; bottom: sensitivity) measure. **C. Incentive effects on confidence formation** Linking incentives, evidence and confidence for correct (left) and incorrect (right) answers. In those two panels, the scatter plots display reported confidence as a function of evidence, for the different incentive levels. The solid line represents the best linear regression fit at the population level. The histograms represent the intercepts (top) and slope for correct (middle) and incorrect answers (bottom) of this relationship, estimated at the individual level and averaged at the population level. The insets presented on the right-hand side of the graph depict the results of the linear mixed-effect model, estimated for each parameter of this regression, i.e., intercept (top), slope for correct answers (middle) and for incorrect answers (bottom). V: net incentive value. |V|: absolute incentive value. Error bars indicate inter-subject standard errors of the mean. *: P<0.05; ** P<0.01; ***P<0.001;

### Effects of incentives on confidence formation

By assuming that confidence builds on noisy perceptual evidence, we expect to observe a positive correlation between confidence and evidence for correct choices and a negative correlation for incorrect choices (see (Fleming and Daw, 2017; Sanders et al., 2016) and **SI results**). Another way of investigating the consequences of confidence incentivization on metacognitive accuracy is to assess how incentives modulate the relationship between confidence and evidence, for correct and incorrect answers: incentive effects can affect confidence per se (suggesting a simple bias of confidence) or influence the relationship between confidence and evidence (suggesting that incentives affect the integration of evidence in the formation of the confidence signal). Although similar in essence to the metacognitive metrics (bias and sensitivity) used above, this approach is model-free, and does not rely on some of the assumptions required for the meta-d’ (Fleming and Lau, 2014; Maniscalco and Lau, 2012). Thus, for each individual and each incentive level, we built a multiple linear regression modeling confidence ratings as a combination of a confidence baseline, and of two terms capturing the linear integration of perceptual evidence for correct and incorrect answers. For each incentive level, regression coefficients were estimated at the individual level and the effect of incentives on the different regression coefficients were subsequently tested in our linear mixed-effect model. Our results show a clear dissociation between the motivational and biasing effects of incentives on confidence formation. On the one hand, the absolute incentive value impacts the slopes of those regressions: in both cases gains and losses increase the linear relationship between confidence and evidence, compared to no incentives (correct answers: β_|V|_ = 0.04 ± 0.02, p < 0.05; incorrect answers: β_|V|_ = -0.24±0.08, p < 0.01; **Fig. 2.C**). On the other hand, the net incentive value affects the intercept of those regressions (β_V_ = 2.18±0.47, p < 0.001; **Fig. 2.C**). This indicates that, while the motivational effect of incentives actually influences the way confidence is built from evidence by increasing the weight of evidence in the ratings in opposite direction for correct and incorrect answers, the biasing effect of incentives appears to be a purely additive effect of incentives on confidence, unrelated to the amount of evidence. These results therefore confirm and extend the reported biasing effects of incentives on overconfidence (metacognitive bias) and the motivational effects of incentives on metacognitive sensitivity.

To further investigate how incentives influence confidence, and to control for alternative explanations, we next conducted three additional experiments.

### The effects of incentives without incentivizing confidence judgements

To rule out that participants deliberately and strategically increase their confidence with net incentive value, due to some misconceptions induced by the incentivization (i.e., MP) mechanisms, we collected data from twenty-one new participants who performed a second task without MP incentivization (i.e. a *performance* task) in addition to our standard *confidence* task. In the performance task, confidence is simply elicited with ratings after choice; confidence accuracy is not incentivized with the MP mechanisms, and subjects are only rewarded according to their choice performance - correct/incorrect (see **Methods** and **Fig. 3**). Still, in line with the value-confidence hypothesis, confidence is found to be specifically modulated by the net incentive value in both the confidence and the performance task (confidence task: β_V_ = 2.47±0.68, p < 0.001; performance task: β_V_ = 1.88±0.59, p < 0.01 **Fig. 4.A**) while incentives have no effects on performance in either task (as expected from the task design; all ps > 0.38).

**Figure 3.**
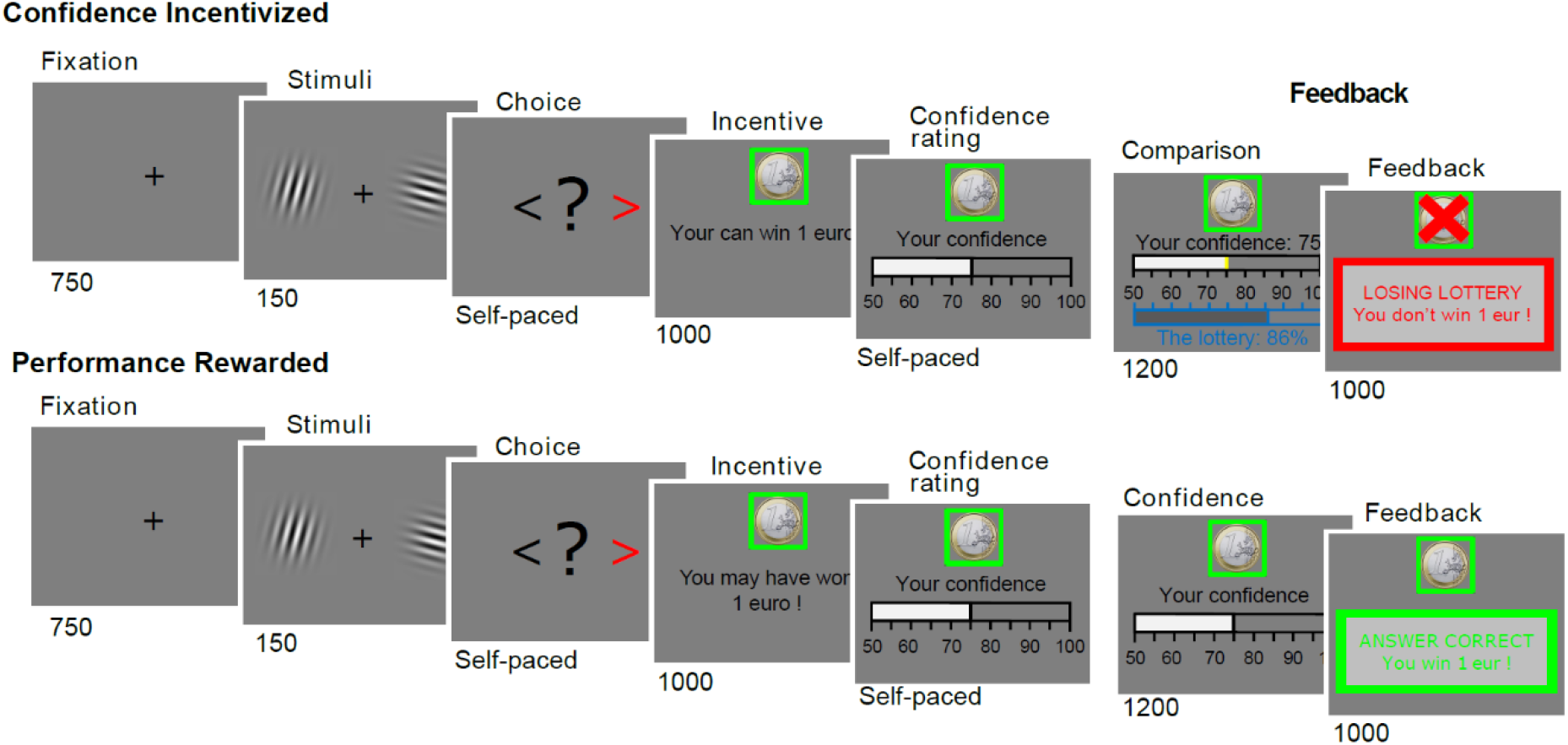
Experiment 2. Successive screens displayed in one trial are shown from left to right with durations in ms. Participants viewed a couple of Gabor patches displayed on both sides of a computer screen, and judged which had the highest contrast. They were thereafter presented with a monetary stake (in a green frame for gain, grey for neutral and red for losses), and asked to report their confidence C in their answer on a scale from 50 to 100%.

**Fig. 4.**
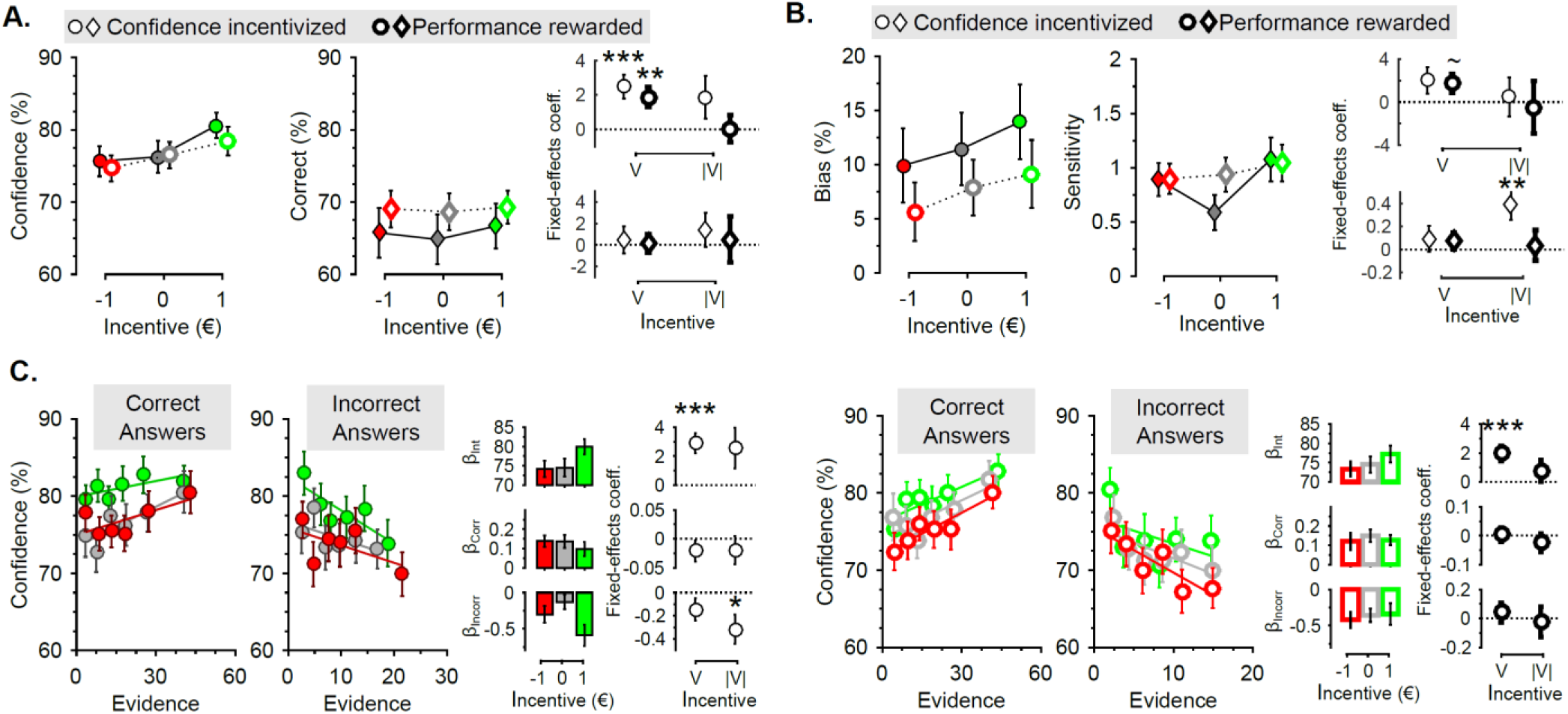
Experiment 2. **A. Incentive effects on behavior.** Reported confidence (dots – leftmost scatter plot) and performance (diamonds – rightmost scatter plot) – i.e. % correct-as a function of incentives. **B. Incentive effects on confidence accuracy.** Computed metacognitive bias (dots – leftmost scatter plots) and sensitivity (diamonds – rightmost scatter plots) as a function of incentives. The insets presented on the right-hand side of the graphs (**A-B**) depict the results of the linear mixed - effect model, estimated for each behavioral (**A**. top: confidence; bottom: performance) and metacognitive (**B.** top: bias; bottom: sensitivity) measure. Empty markers with thick edges indicate the Performance Rewarded task. **C. Incentive effects on confidence formation** Linking incentives, evidence and confidence for the Confidence incentivized (left half) and the Performance Rewarded (right half) tasks, for both correct (left scatterplot) and incorrect (right scatterplot) answers. In those two panels, the scatter plots display reported confidence as a function of evidence, for the different incentive levels. The solid line represents the best linear regression fit at the population level. The histograms represent the intercepts (top) and slope for correct (middle) and incorrect answers (bottom) of this relationship, estimated at the individual level and averaged at the population level. The insets presented on the right-hand side of the graph depict the results of the linear mixed-effect model, estimated for each parameter of this regression, i.e., intercept (top), slope for correct answers (middle) and for incorrect answers (bottom). V: net incentive value. |V|: absolute incentive value. Error bars indicate inter-subject standard errors of the mean. *: P<0.05; ** P<0.01; ***P<0.001;

Regarding confidence accuracy, the effect of the net incentive value percolates metacognitive bias in both tasks, although merely as a trend (confidence task: β_V_ = 2.01±1.24, p = 0.11; performance task: β_V_ = 1.75±0.89, p = 0.05; **Fig. 4.B**). Interestingly, the motivational effect of the absolute incentive value on metacognitive sensitivity is replicated when confidence is incentivized, but not when performance is incentivized (confidence task: β_|V|_ = 0.40 ± 0.14, p < 0.01; performance task: β_|V|_ = 0.04 ± 0.13, P = 0.79; **Fig. 4.B**). We then replicate and extend the findings of the first experiment on the confidence formation model (**Figure 4.C**): when biasing effects of the net incentive value are present (in both tasks), they impact the intercept of the confidence formation model (both ps < .001). On the other hand, the motivational effects of the absolute incentive value are only found on the slope of incorrect trials in the confidence task (incorrect answers; confidence task: β_|V|_ = -0.317±0.13, p < 0.05; performance task: β_|V|_ = -0.03±0.10, p = 0.81). Again, this means that the motivational effect of the incentivization of confidence accuracy is underpinned by a better integration of perceptual evidence in the confidence rating when stakes increase, whereas this effect is absent in the task where confidence accuracy is not incentivized. In sum, these results indicate that the biasing effects of incentives on confidence judgments are not induced by the incentivization mechanism, and that the motivational effects of incentives on confidence and metacognitive accuracy are only found when confidence is incentivized.

### Dissociating incentive value effects from (mere) valence effects

In order to demonstrate that the motivational and biasing effects of incentives are due to incentive *values*, rather than to simple valence (gain/loss) effects, we next invited thirty-five subjects to participate in a third experiment, where incentives for confidence accuracy varied in both valence (gains and losses) and magnitude (1€ vs 10¢) (see **Table 1**). We modified our linear mixed-effect models to include a valence variable (=1 if incentives are positive and 0 if negative, thereafter indexed by +/-), in addition to the net and absolute incentive value. Results show that both the net incentive value and the valence variable impact confidence judgments (β_V_ = 1.02 ± 0.38, p < 0.01; β+/- = 4.01±1.11, p < 0.001; **Fig. 5.A**). This means that the biasing effects of incentive previously reported are not simply due to an effect of valence, but are truly underpinned by the net incentive value. Again, as designed and expected, no effects of incentives are found on performance (all ps > 0.22). Importantly, the linear effect of the net incentive value transfers to the metacognitive bias (β_V_ = 2.15 ± 0.97, p < 0.05; **Fig. 5.B**). Note that we do not find significant effects of the absolute value of incentives on metacognitive sensitivity (all ps > 0.25 **Fig. 5.B**). This difference with the results of the two previous experiments can be explained by the lack of a neutral incentive condition in the present experiments. This means that motivational effects previously reported would be primarily due to the mere presence of incentives.

**Fig. 5.**
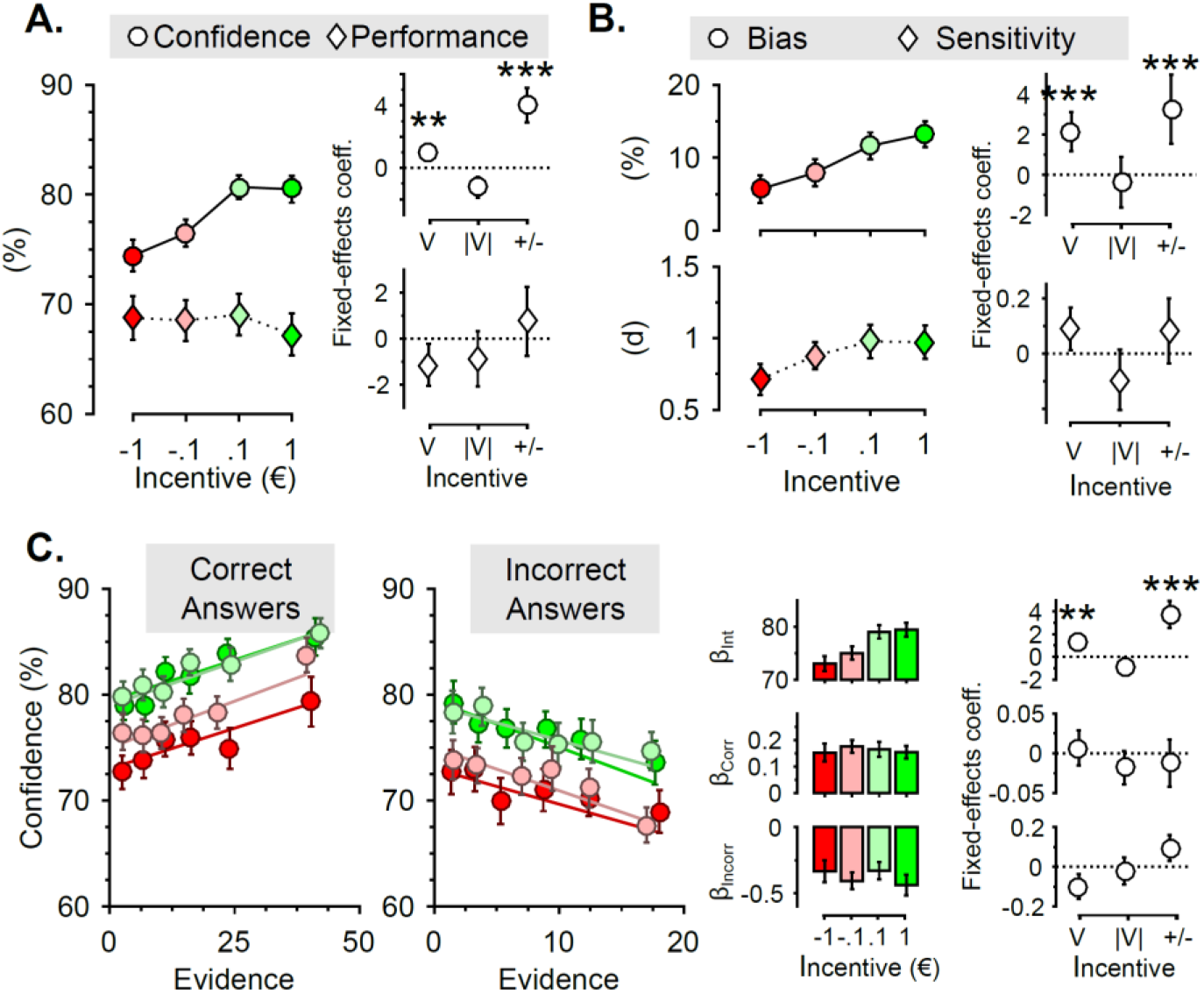
Experiment 3. **A. Incentive effects on behavior.** Reported confidence (dots) and performance (diamonds) – i.e. % correct-as a function of incentives. **B. Incentive effects on confidence (metacognitive) accuracy.** Computed bias (top, dots) and meta-d’ (diamonds) as a function of incentives. The insets presented on the right-hand side of the graphs (**A-B**) depict the results of the linear mixed-effect model, estimated for each behavioral (**A**. top: confidence; bottom: performance) and metacognitive (**B.** top: bias; bottom: sensitivity) measure. **C. Incentive effects on confidence formation** Linking incentives, evidence and confidence for correct (left) and incorrect (right) answers. In those two panels, the scatter plots display reported confidence as a function of evidence, for the different incentive levels. The solid line represents the best linear regression fit at the population level. The histograms represent the intercepts (top) and slope for correct (middle) and incorrect answers (bottom) of this relationship, estimated at the individual level and averaged at the population level. The insets presented on the right-hand side of the graph depict the results of the linear mixed-effect model, estimated for each parameter of this regression, i.e., intercept (top), slope for correct answers (middle) and for incorrect answers (bottom). V: net incentive value. |V|: absolute incentive value. +/-: incentive valence. Error bars indicate intersubject standard errors of the mean. *: P<0.05; ** P<0.01; ***P<0.001;

Replicating our previous finding, the biasing effect of the net incentive value is found to be independent from the amount of evidence, impacting the intercepts of the linear relationship between evidence and confidence (intercept: β_V_ = 1.34±0.50, p < 0.01; **Fig. 5.C**). No effects of incentives could be found on the slopes characterizing the integration of evidence in confidence judgments (all ps > 0.11). In sum, the results from this third experiment replicate the biasing effect of the net incentive value on confidence, and furthermore demonstrate that these effects depend on the magnitude of incentives.

### Accounting for difference between gain and loss in effect on confidence

While the effect of the net incentive value on confidence and metacognitive bias revealed in our first three experiments appeared robust and replicable, it seemed to be driven by the loss frame. This could mean that this biasing effect is purely restricted to the loss frame. However, an alternative hypothesis is that subjects are simply less sensitive to gains, as suggested by prospect theory (Tversky and Kahneman, 1991). In order to dissociate between those two hypotheses, we invited twenty-four subjects to participate in a final study which included higher stakes (10c, 1€, 2€) in both gain and loss frames (**Table 1**). In this case, our linear mixed-effect model included three independent variables: two variables accounted for the signed incentive magnitude in the gain (V+), and in the loss frame (V-); in addition, and in line with the previous experiment, the third variable captured the effect of the valence framing (+/-). Our results reveal a significant effect of the incentive magnitude on confidence, in both the gain and loss frames (β_V+_ = 20.79±0.25, p < 0.001; β_V-_ = 1.22±0.38, p < 0.001; **Fig. 6.A**), and no effects of incentives on performance (all ps > 0.45). This result confirms our initial hypothesis: following expected values, higher incentives seem to bias confidence judgments upwards in a gain frame, and downward in a loss frame.

**Fig. 5.**
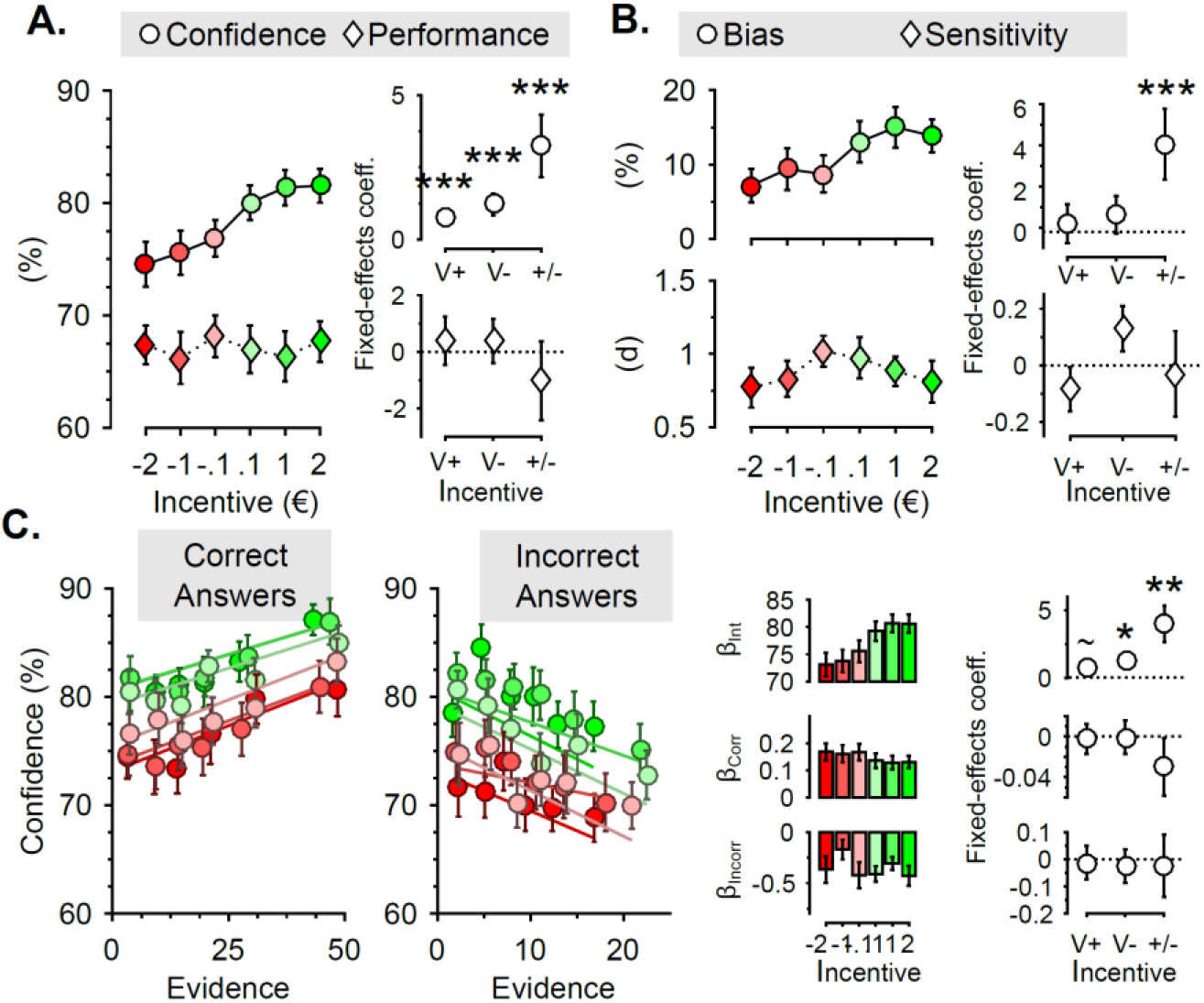
Experiment 4. **A. Incentive effects on behavior.** Reported confidence (dots) and performance (diamonds) – i.e. % correct-as a function of incentives. **B. Incentive effects on confidence (metacognitive) accuracy.** Computed bias (top, dots) and meta-d’ (diamonds) as a function of incentives. The insets presented on the right-hand side of the graphs (**A-B**) depict the results of the linear mixed-effect model, estimated for each behavioral (**A**. top: confidence; bottom: performance) and metacognitive (**B.** top: bias; bottom: sensitivity) measure. **C. Incentive effects on confidence formation** Linking incentives, evidence and confidence for correct (left) and incorrect (right) answers. In those two panels, the scatter plots display reported confidence as a function of evidence, for the different incentive levels. The solid line represents the best linear regression fit at the population level. The histograms represent the intercepts (top) and slope for correct (middle) and incorrect answers (bottom) of this relationship, estimated at the individual level and averaged at the population level. The insets presented on the right-hand side of the graph depict the results of the linear mixed-effect model, estimated for each parameter of this regression, i.e., intercept (top), slope for correct answers (middle) and for incorrect answers (bottom). V+: net incentive value for gains. V-: net incentive value for losses. +/-: incentive valence. Error bars indicate inter-subject standard errors of the mean. ~P<0.10; *: P<0.05; ** P<0.01; ***P<0.001;

Similar to our third experiment, no motivational effect of incentives is detectable on metacognitive sensitivity (all ps > 0.11; **Fig. 6.B**), suggesting that it is mostly driven by the incentive versus no-incentive contrast. Slightly departing from what is observed in the previous experiments, the effects of incentives on metacognitive bias are, this time, mostly driven by the valence variable (β_+/-_ = 4.26±1.72, p < 0.05; **Fig. 6.B**). Given that this measure combines the confidence and performance variance, and that the presence of six incentive-levels decreases the number of trials used to estimate those measure, we interpret the absence of effects of the magnitude of incentives on metacognitive bias as a lack of power. Supporting this interpretation, the biasing effects can be found on the intercept of our confidence formation model, a more sensitive measure of our bias (β_V+_ = 0.72±0.40, p = 0.08; β_V-_ = 1.26±0.58, p < 0.05; **Fig. 6.C**). This last set of results replicates, for the fourth time, the biasing effects of incentives on confidence, and confirms that monetary gains and losses both contribute to biasing confidence in perceptual decisions.

## Discussion

In this article, we combined a perceptual decision task and an auction procedure inspired from behavioral economics (Becker et al., 1964; Ducharme and Donnell, 1973) to investigate how monetary incentives influence confidence. In addition to replicating important features of a recent model of confidence formation (Sanders et al., 2016), we reveal and dissociate two effects of monetary incentives on confidence accuracy.

The first effect is a motivational effect of incentives: in line with theories of rational decision making and motivation, incentivizing confidence judgments improves meta-cognitive sensitivity. This means that high (respectively low) confidence is more closely associated with correct (respectively incorrect) decisions when confidence reports are incentivized, regardless of the valence or magnitude of the incentive. This nicely extends a recent study reporting a similar effect of incentivization on discrimination (a measure closely related to sensitivity, assessing how confidence discriminate between correct and incorrect answers), although limited to the gain domain (Hollard et al., 2015). This also confirms that the MP mechanisms is particularly well-suited to investigate confidence incentivization (Hollard et al., 2015; Massoni et al., 2014). Here, we further show that this motivational effect of incentives is underpinned by a better integration of perceptual evidence in the confidence judgment when stakes increase.

The second effect, the biasing effect of incentives, is more striking: confidence judgments are parametrically biased by the net value of the incentive. The prospect of gains increases confidence, while the prospect of losses decreases confidence. Because people generally exhibited overconfidence in our experiment, the gain prospects detrimentally increased the overconfidence bias, while prospects of losses reduced this bias and improved confidence accuracy. As opposed to the motivational effect, the biasing effect of incentive was purely additive, i.e., independent of the amount of evidence on which decisions and confidence judgments are based. The biasing effect was also found to be incidental, i.e., also present when the confidence judgments are not incentivized. Importantly, we show that this bias is unpredicted by motivated cognition theories such as the desirability bias (Giardini et al., 2008), which predicts that the overconfidence bias would also increase with negative incentive values, because avoiding a loss is desirable. This biasing effect is also unpredicted by the theories of rational decision making and motivation, which predicts decreased overconfidence with increased positive incentive values because it would lead to a higher reward (as incentivized by the MP mechanism). Yet, the biasing effect of incentives is in line with the value-confidence hypothesis. One plausible interpretation for this effect is an affect-as-information effect: people use their momentary affective states as information in decision-making (Schwarz and Clore, 1983) which, in our case, means that they integrate the trial expected value into their confidence judgment. These results and interpretations nicely fit with recent reports showing that negative affective states (such as worry) decrease overconfidence (Massoni, 2014), while positive affective states (such as joy) increase overconfidence (Koellinger and Treffers, 2015). The reported effects of incentive on confidence also confirm that confidence judgments do not just represent rational estimates of the probability of being correct (Pouget et al., 2016), but additionally integrate information and potential biases processed after a decision is made (Fleming and Daw, 2017; Navajas et al., 2016). These results therefore provide additional evidence in favor of second-order models of confidence, which propose that confidence builds on different samples of evidence than the ones used to render the decision (Fleming and Daw, 2017).

In order to incentivize confidence reports, we used a mechanism inspired from Becker-DeGroot-Marschak auction procedures (Becker et al., 1964; Ducharme and Donnell, 1973), referred to as Reservation or Matching Probability (MP), which conveniently allowed us to manipulate the monetary stakes on a trial-by-trial basis. Contrary to other incentivization methods such as the quadratic scoring rule (QSR), the MP mechanism is valid under simple utility maximization assumptions, i.e., remains incentive-compatible when subjects are not risk neural (Hollard et al., 2015; Schlag et al., 2015). The MP mechanism is even incentive-compatible when considering probability distortions, on the assumption that both subjective (confidence) and objective (lotteries) probabilities are transformed identically (Trautmann and van de Kuilen, 2015; Wakker, 2004). Several studies have investigated the impact of different incentivization mechanisms on subjective probability judgments (confidence or belief), and report that MP is among the best methods available, both at the theoretical and experimental levels (Hollard et al., 2015; Schlag et al., 2015; Trautmann and van de Kuilen, 2015) and is particularly well-suited for signal-detection theory analyses (Massoni et al., 2014). Our findings nonetheless demonstrate that regardless of the type of incentivization mechanism, the mere presence of incentives can induce distortions in probability estimations such as confidence judgments.

In this collection of experiments, we only used relatively small monetary amounts as incentives; how the motivational and biasing effects of incentives scale when monetary stakes increase significantly remains an open question. Critically, higher stakes may also impact physiological arousal, which influence confidence and interoceptive abilities (Allen et al., 2016; Kandasamy et al., 2016). In general, the effects of incentives on confidence accuracy could also be mediated by inter-individual differences in metacognitive or interoceptive abilities (Kandasamy et al., 2016; Song et al., 2011) and by incentive motivation sensitivity (Harsay et al., 2011). Because our subject sample was mostly composed of university students, the generalization of those findings in the general population will have to be assessed in further studies.

The mere notion of confidence biases, notably overconfidence, and the actual conditions under which they can be observed sparked an intense debate in psychophysics (Baranski and Petrusic, 1994; Juslin et al., 2000; Olsson and Winman, 1996) and evolutionary theories (Johnson and Fowler, 2011; Marshall et al., 2013). Critically, here, confidence accuracy was properly incentivized, hence deviations from perfect calibration can be appropriately interpreted as cognitive biases (Marshall et al., 2013). The striking effects of net incentive values on confidence seem to make sense when considering evolutionary perspective: in natural settings, whereas overconfidence might pay off when prospects are potential gains (e.g., when claiming resources (Johnson and Fowler, 2011)), a better calibration might be more appropriate when facing prospects of losses (e.g., death or severe injuries), given their potential dramatic consequences on reproductive chances. Interestingly, the observed valence difference in the effect of incentives magnitude – higher in the loss than in the gain domain-seem to mimic valence asymmetries observed in economic decision-making theories such as prospect theory (Tversky and Kahneman, 1991).

How confidence is formed in the human brain and how neurophysiological constraints explain biases in confidence judgments remain an open question (Meyniel et al., 2015b; Pouget et al., 2016). Although functional and structural neuroimaging studies initially linked confidence and metacognitive abilities to dorsal prefrontal regions (Fleming and Dolan, 2012), confidence activations were also recently reported in the ventro-medial prefrontal cortex (De Martino et al., 2013; Lebreton et al., 2015), and in striatal and mesolimbic regions (Guggenmos et al., 2016; Hebart et al., 2014). This network has been consistently involved in motivation and value-based decision making (Levy and Glimcher, 2012). It is therefore possible that this network plays a role in the motivational and biasing effects of incentives on confidence. However, this remains highly speculative and should be investigated in future neuroimaging studies. Overall, our results suggest that investigating the interactions between incentive motivation and confidence judgments might provide valuable insights on the cause of confidence miscalibration in healthy and pathological settings. For instance, high monetary incentives in financial or managerial domains may in fact create or exaggerate overconfidence, leading to overly risky and suboptimal decisions. In the clinical context, inflated levels of overconfidence in pathological gamblers (Goodie, 2005) could be amplified by high monetary incentives, contributing to compulsive gambling in the face of great loss. Moreover, if indeed value-induced affective states modulate confidence judgements, other disorders with abnormal incentive processing such as addictions, mood-disorders, obsessive-compulsive disorder and schizophrenia could be at particular risk for confidence miscalibration (Figee et al., 2011; Goldstein et al., 2007; Whitton et al., 2015). Field experiments and clinical research will be needed to further explore the individual and societal consequences of the interactions between incentive motivation and confidence accuracy.

## Experimental Procedures

### Subjects

All studies were approved by the local Ethics Committee of the University of Amsterdam Psychology Department. All subjects gave informed consent prior to partaking in the study. The subjects were recruited from the laboratory's participant database (https://www.lab.uva.nl/spt/). A total of 83 subjects took part in this study (see Table 1). They were compensated with a combination of a base amount (10€), and additional gains and/or losses from randomly selected trials (one per incentive condition per session for experiment 1, and one per incentive condition from one randomly selected session for experiments 2 and 3).

### Tasks

All tasks were implemented using MATLAB^®^ (MathWorks) and the COGENT toolbox (http://www.vislab.ucl.ac.uk/cogent.php). In all four experiments, trials of the confidence incentivization task shared the same basic steps (**Fig. 1.A**): after a brief fixation cross (750 ms), participants viewed a pair of Gabor patches displayed on both sides of a computer screen (150 ms), and judged which had the highest contrast (self-paced) by using the left or right arrow. They were thereafter presented with a monetary stake (1000 ms, accompanied by the sentence “You can win[/lose] X euros”) and asked to report their confidence C in their answer on a scale from 50 to 100% by moving a cursor with the left and right arrows, and selecting their desired answer by pressing the spacebar (self-paced). The initial position of the cursor was randomized between 65 and 85% to avoid anchoring of answers on 75%. The steps following the confidence rating, and the relation between the monetary stake, the confidence and the correctness of the answer were manipulated in two main versions of this task. In the ***Extended Version,*** at the trial level, the *lottery draw* step was separated in two smaller steps. First, a lottery number L was drawn in a uniform distribution between 50 and 100% and displayed as a scale under the confidence scale. After 1200 ms, the scale with the highest number was highlighted for 1200 ms. Then, during the *resolution step*, if C happened to be higher than L, a clock was displayed for 750 ms together with the message “*Please wait*”. Then, a *feedback* was displayed which depended on the correctness of the initial choice. Back at the *resolution step*, if the L happened to be higher than C, the lottery was implemented. A wheel of fortune, with a L% chance of losing was displayed, and played; the lottery arm spun for ~750 ms, and would end up in the winning (green) area with L% probability or in the losing (red) area with 1-L% probability. Then, a *feedback* informed whether the lottery was winning or losing.

Subject would win (gain frame) or not lose (loss frame) the incentive in case of a “winning” trial, and they would not win (gain frame) or lose (loss frame) the incentive in case of a “losing” trial. Thanks to the MP procedure, the strategy to maximize one's earnings is to always report on the confidence scale one's subjective probability of being correct as truthfully and accurately as possible (**SI Text**).

Subjects were explicitly instructed so. In addition to extensive instructions explaining the MP procedure, participants gained direct experience with this procedure through a series of 24 training trials that did not count towards final payment.

In the ***Short version***, the incentivization scheme was the same as in the ***Extended Version***, but part of it was run in the background. Basically, the lottery scale appeared, and the scale with the highest number was highlighted concomitantly (1200ms). Additionally, the *resolution step* was omitted. Still, the complete feedback relative to the lottery and or the correctness of the answer was given to subjects in the *feedback* step. There was no difference in our participants' behavior when using the extended or short version of our task.

In the ***Performance Version*** the MP mechanism was omitted, but the layout was similar to the ***Short Version*** (see **Sup. Fig. 2**). The monetary stake screen was accompanied by a different sentence (You may have won[/lost] X euros). The *lottery draw* /comparison step was replaced with a screen of similar duration (1200ms) simply displaying the confidence scale and the chosen rating. A *feedback*, on the correctness of the answer and the trial outcome was finally given at every trial (1000ms).

### Stimuli & design

Participants initially performed a 144 trials calibration session (~5min), where they only performed the Gabor contrast discrimination task, without incentive or confidence measure (**Fig. 1.A**). During this calibration, the distribution of contrast difference (i.e. difficulty) was adapted every 12 trials following a staircase procedure, such that performance reached approximatively 70% correct. The calibration data was used to estimate individual psychometric function:

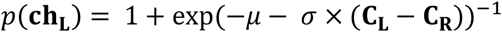

where *p*(**ch_L_**) is the probability of subjects choosing the left Gabor, **C_L_** and **C_R_** and are the contrast intensities of the left and right Gabors. In this formalization, μ quantifies subjects' bias toward choosing the left Gabor in the absence of evidence, and σ quantifies subjects' sensitivity to contrast difference. The estimated parameters (μ and σ) were used to generate stimuli for the confidence task, spanning defined difficulty levels (i.e. known *p*(**ch_L_**)) for all incentives levels. After the first session of the confidence task, μ and σ were re-estimated for each session from the data of the preceding session (experiments 1, 2 and 4), or from a new calibration session (experiment 3).

## Metacognitive metrics

We used two components of metacognition: metacognitive bias and metacognitive sensitivity. Metacognitive bias is obtained by computing the difference between the mean confidence and the mean accuracy.

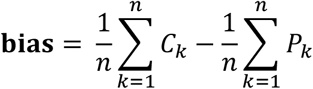

where n is the total number of trials, Ck is the reported confidence at trial k, and Pk is the performance at trial k (1 for a correct answer and 0 for an incorrect answer);

Metacognitive sensitivity was measured as the meta-d’, a new metric introduced by Maniscalco and Lau (2012). Meta-d’ defines the level of d’ that an SDT-ideal observer would need to generate an observed set of confidence ratings, given an observed set of choices. Meta-d’ was computed using the Matlab code of Maniscalco and Lau (2012) available on their website^1^.

Note that all results obtained with the meta-d’ were also replicated with a very simple (but not bias-free) metric of sensitivity, computed as the difference between the average confidence for correct answers and the average confidence for incorrect answers.

### Statistics

All statistical analyses were performed with Matlab R2015a. All statistical analyses reported in the main text result from linear mixed-models (estimated with the glmefit function). For each (non-reaction time) behavioral (e.g. confidence, performance) and metacognitive (bias, sensitivity) measure Y, we computed the average of Y per incentive level per individual. For reaction-times, whose distribution are typically skewed, we computed the median, rather than the mean reaction time in each incentive condition. For the confidence formation model, we used the regression coefficient from the individual linear regressions linking confidence and evidence for correct and incorrect choices, estimated per individual and incentive level. We then used as predictor variables the absolute incentive value (|V|), the net incentive value (V) and the incentive valence (+/-, only for Experiments 3 and 4). All mixed models included random intercepts and random slopes. As an example, in Wilkinson-Rogers notation, the linear mixed-effect models for Experiment 1 can be written: Y ~ 1 + |V| + V + (1 + |V| + V |Subject). Detailed results on all linear mixed-effect models used in the manuscript can be found in the **SI results**.

## Acknowledgments

ML, JL and RJvH were supported by Amsterdam Brain and Cognition (ABC) Talent Grants (Universiteit van Amsterdam). ML was additionally supported by an NWO Veni Fellowship, and the Bettencourt Schueller Fondation. AEG was supported by an NWO VIDI grant. The authors thank Stefano Palminteri, Ivan Soraperra, Frans van Winden, Jan B. Engelmann and Joel van der Weele for helpful discussions and comments on the manuscript, and Thomas A. Davison for checking the English.

## Footnotes

1 (http://www.columbia.edu/~bsm2105/type2sdt/)

